# Human Reference Genome and a High Contiguity Ethnic Genome AK1

**DOI:** 10.1101/795807

**Authors:** Jina Kim, Joohon Sung, Kyudong Han, Wooseok Lee, Seyoung Mun, Jooyeon Lee, Kunhyung Bahk, Inchul Yang, Young-Kyung Bae, Changhoon Kim, Jong-il Kim, Jeongsun Seo

## Abstract

Studies have shown that the current human reference genome (GRCh38) might miss information for some populations, but “exactly what we miss” is still elusive due to the lower contiguity of non-reference genomes. We juxtaposed the GRCh38 with high contiguity genome assemblies, AK1, to show that ∼1.8% (∼53.4 Mbp) of AK1 sequences missed in GRCh38 with ∼0.76% (∼22.2 Mbp) of ectopic chromosomes. The unique AK1 sequences harbored ∼1,390 putative coding elements. We found that ∼5.3Mb (∼0.2%) of the AK1 sequences aligned and recovered the “unmapped” reads of fourteen individuals (5 East-Asians, 4 Europeans, and 5 Africans) as a reference. The regions that “unmapped” reads aligned included 110 common (shared between ≥2 individuals) and 38 globally (≥7 individuals) missing regions with 25 candidate coding elements. We verified that many of the common missing regions exist in multiple populations and chimpanzee’s DNA. Our study illuminates not only the discovery of missing information but the use of highly precise ethnic genomes in understanding human genetics.

## INTRODUCTION

DNA Sequencing of the human genome is the pivot of precision medicine. For the sake of economy, an overwhelming majority of sequencing platforms adopt re-sequencing methods, using the GRCh37/38 (a.k.a., hg19/38) human genome assembly as the reference. The GRCh38 is the heir of the Human Genome Project and has been further enriched (∼30%) by the addition of >50 individuals’ genome, including African Ancestry(Schneider et al. 2017). It has been generally believed that a single global reference genome would suffice because the re-sequencing requires a reference to determine individuals’ genetic variants rather than a database encompassing a list of variants. However, recent findings that ethnic diversities in the structural variations are substantial (Genomes Project et al. 2015; Sudmant et al. 2015), and that the human migration has complex detour and local admixture(Mondal et al. 2016) ensued questions whether some part of DNA sequences are missing by current re-sequencing practice (Maretty et al. 2017).

The “unmapped” are sequence reads that failed to align the reference. Previous studies examined the unmapped reads to identify regions with suggestive evidence of protein-coding (Sherman et al. 2018) or diseases association (Kehr et al. 2017; Maretty et al. 2017). Studies used fragmented raw genome data, and performed *de novo* assembly of different individuals’ sequences in order to compare with the reference(Maretty et al. 2017; Sherman et al. 2018; Duan et al. 2019). This approach, however, lacks chromosomal landscape and proportionality, and admixes “missing” regions with “unmappable” sequences primarily due to low contiguity of ethnic genome data. For example, the approximated missing genome of ∼300Mb for Africans through a *de novo* assembly of the unmapped reads(Sherman et al. 2018) might have been over-estimated by the chimeric regions. In this study, we performed an exact comparison of the two most contiguous human genome assemblies, the GRCh38 and AK1(Seo et al. 2016). Additionally, we found the existence and putative function of commonly missing parts and verified the presence of missing regions with high contiguity ethnic genome.

## RESULTS

### Systematic comparison between GRCh38p12 and AK1

We first compared the whole AK1 sequences on the GRCh38 (“liftover”) and its alternative sequences, to search for synteny. By generating bi-directional “chain files” indicating both homology and gaps of base-pair resolution, we categorized a total of 2,832 scaffolds (∼2.90 Gbp) of AK1 into three by the alignment patterns: the first scaffolds group (n=945, sum of ∼2.70Gbp) consists ≥ 99% of chromosomes of the GRCh38; the third scaffold group (n=1,420 ∼41Mbp) lacked synteny with the GRCh38; the second group (n=467, ∼165Mbp) had partial (<99%) matches. A total of 53.4 Mbp (∼1.8%) of the AK1 genome lacked homology with GRCh38. A total of 899 AK1 scaffolds were hybrids (Group 1 and 2) of multiple chromosomes of the GRCh, in which compound, ectopic chromosomes’ contribution amounted to ∼22.2 Mbp (∼0.76 %) (Fig. 1).

**Figure 1.**
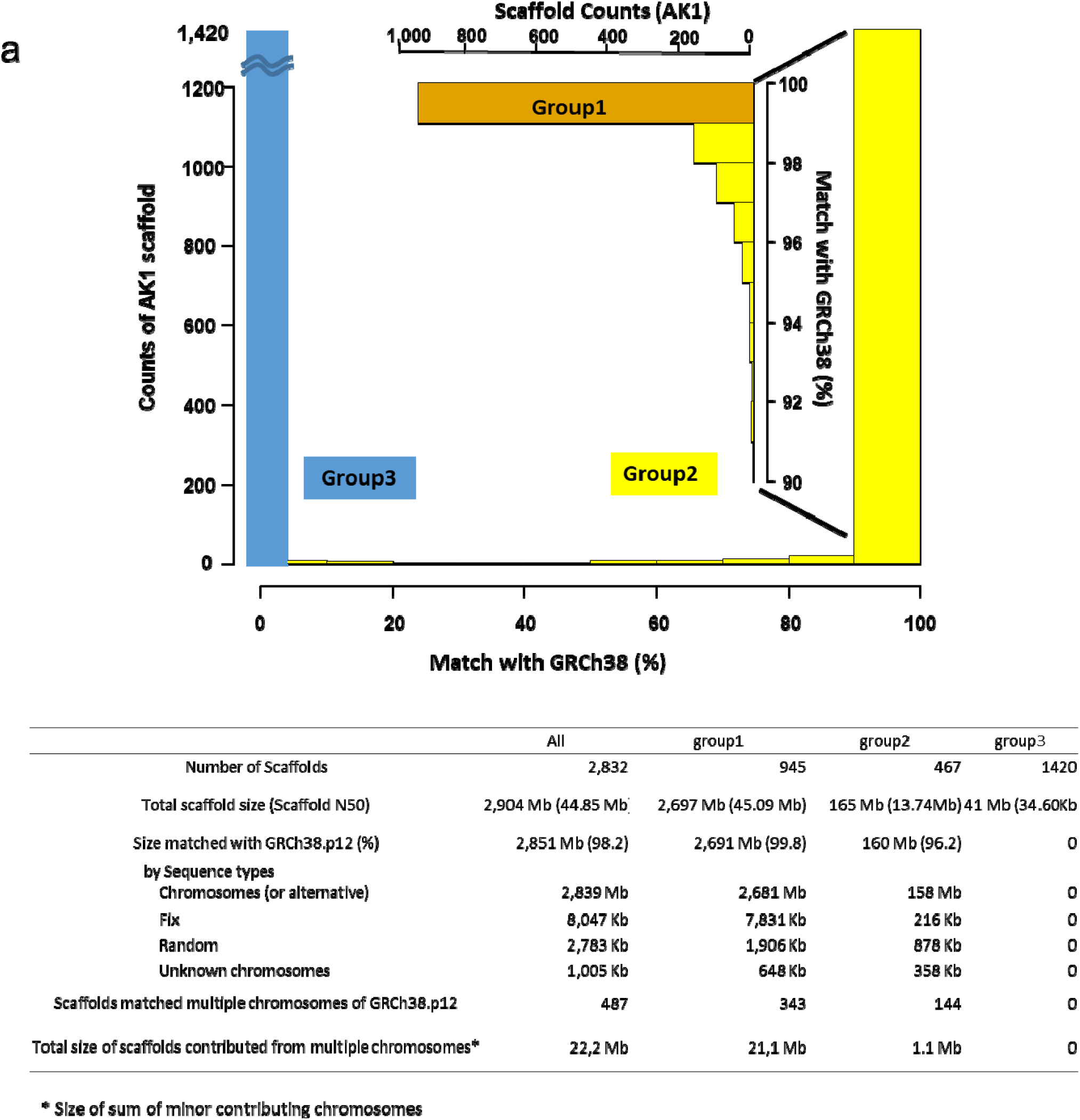
A systematic comparison between AK1 scaffolds (n=2,382) and GRCh38.p12. The degree of match divided AK1 scaffolds into three distinct patterns of synteny by LASTZ(Harris. 2007). (a) The x axis (and vertical popup axis for the group 1) represents percent of matches between AK1 scaffold and GRCh38.p12 chromosomes, and the y axis represents count of scaffolds (b) Statistics of three groups of AK1 scaffolds according to the above matching patterns.

For finding the genome sequences unique to AK1, we selected 3,333 regions larger than 200bp and searched for protein-coding function through a translated BLAST(Camacho et al. 2009) within mammals. A total of 1390 regions (e-value <10^−10^, identity ≥ 70%, and alignment length ≥ 50bp) were predicted to have putative protein-coding elements (Supplemental Table S1). The group 3 scaffolds, which are unique to AK1, had different components of genome sequences and repeats when we analyzed using the RepeatMasker(Smit 2015). Satellite repeat sequences preponderated (> 87%) with a higher proportion of simple repeats (Supplemental Table S2)

### Profile of the “Unmapped Reads”

We selected 14 individuals’ in-depth (>50X) WGS data from the 1kG database comprising Caucasians (4 individuals), Asians (5 individuals), and African ancestry (5 individuals). All data represented different area of populations, and were initially aligned against GRCh38 with written information of the quality control (all by the Illumina HiSeq platform). In average, ∼4.7% of the WGS data (∼2.6 M out of 54.6M total reads per individual) failed to align GRCh38 including its alternative sequences. Africans had the lowest, and Caucasians had the highest mapping rate on GRCh38 (Supplemental Table S3). “Unpaired reads” explained the most substantial part of the unmapped (∼59%) (Supplemental Fig. S1), due to the differences in sequencing quality between read1 and read2 (both ends of complimentary sequences). Besides the generally lower sequencing quality, the proportion of repetitive sequences among unmapped reads was about ten times more low-complexity and >2 times more simple repeats and satellites, whereas they had much lower proportion of SINE, LINE, and LTR compared with reference genome (Supplemental Table S4). The use of massive fragmented reads will inevitably generate equivocal data for alignment, particularly on satellite or low complexity regions. Given the quality and components, technical characteristics intrinsic to the analytic platform rather than the incompleteness of the reference genome, is likely to spawn the majority of the unmapped reads.

### Recovered genomic regions by the “Re-alignnent” to AK1

In average, 72K out of ∼2.6 M reads per individual (mapping quality >10) were newly mapped on AK1, with a tiny proportion of microbial origins (∼0.01%) (Table1). The recovery rates by the re-alignment to AK1 were relatively low (overall 0.92% or 0.49% for high fidelity mapping quality) and did not show substantial differences between populations (Table 1). The recovered region amounted to ∼0.2% (5.3Mb) of AK1 sequences (Fig 2b). We drilled down the recovered sequences into three groups of the Fig1 (Fig. 2b). Group1 scaffolds harbored the largest number (n=58,340) of the re-aligned reads, but proportion-wise, Group3 scaffolds populated unmapped reads more widely. For the regions on Group1 (≥ 99% homology with GRCh38), most of the alignment arose on the putative insertions, absent in GRCh38 sequences (Fig. 2c). Regions in homology with chromosome 19 and 21 were most densely mapped (Supplemental Fig.S2 and Supplemental Table S5). Our findings suggest that the addition of ethnic reference do salvage missing genome regions, although a small part of the “unmapped reads” gave the results.

**Table 1.**
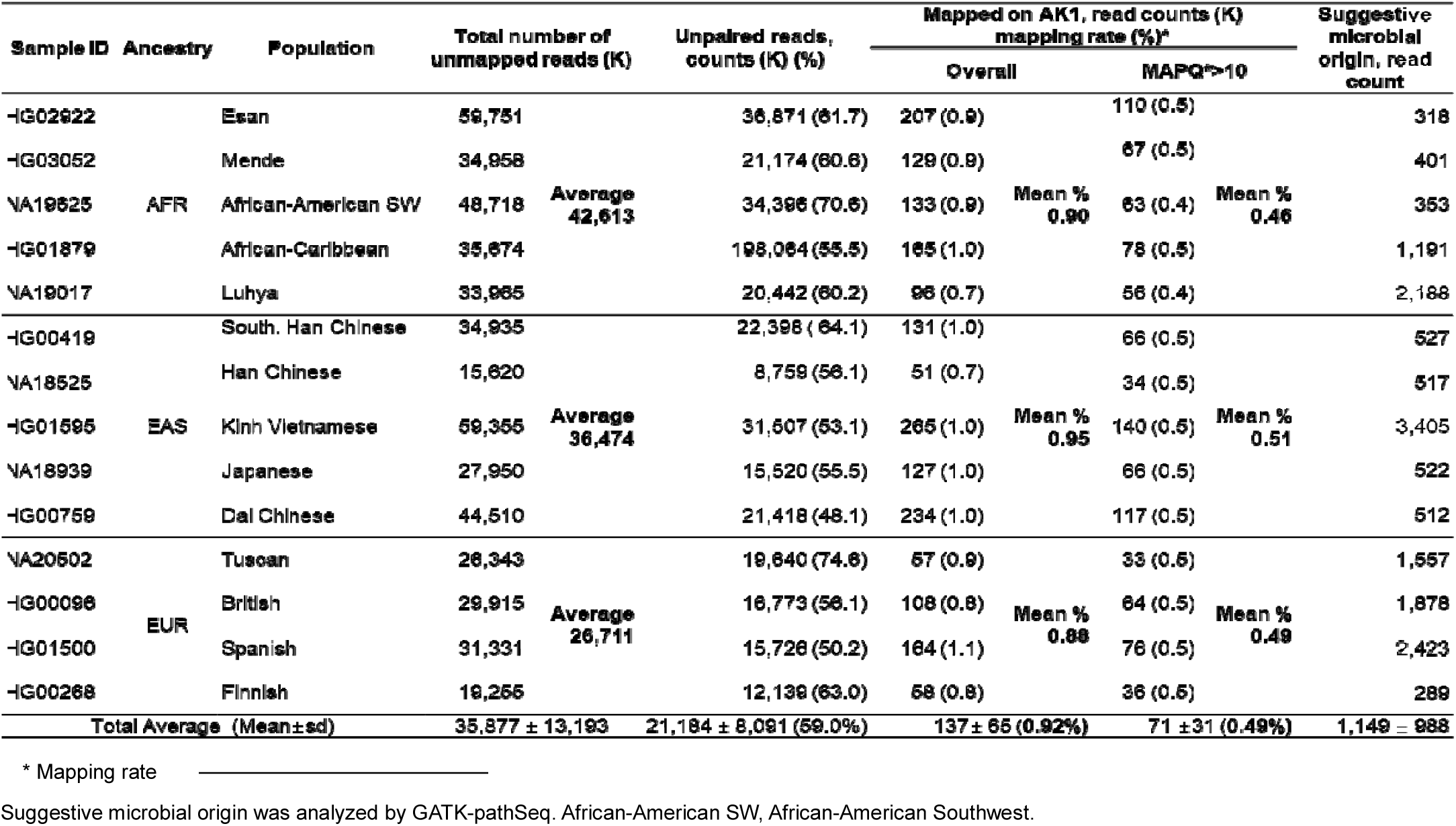
The read counts of unmapped reads by samples

**Figure 2.**
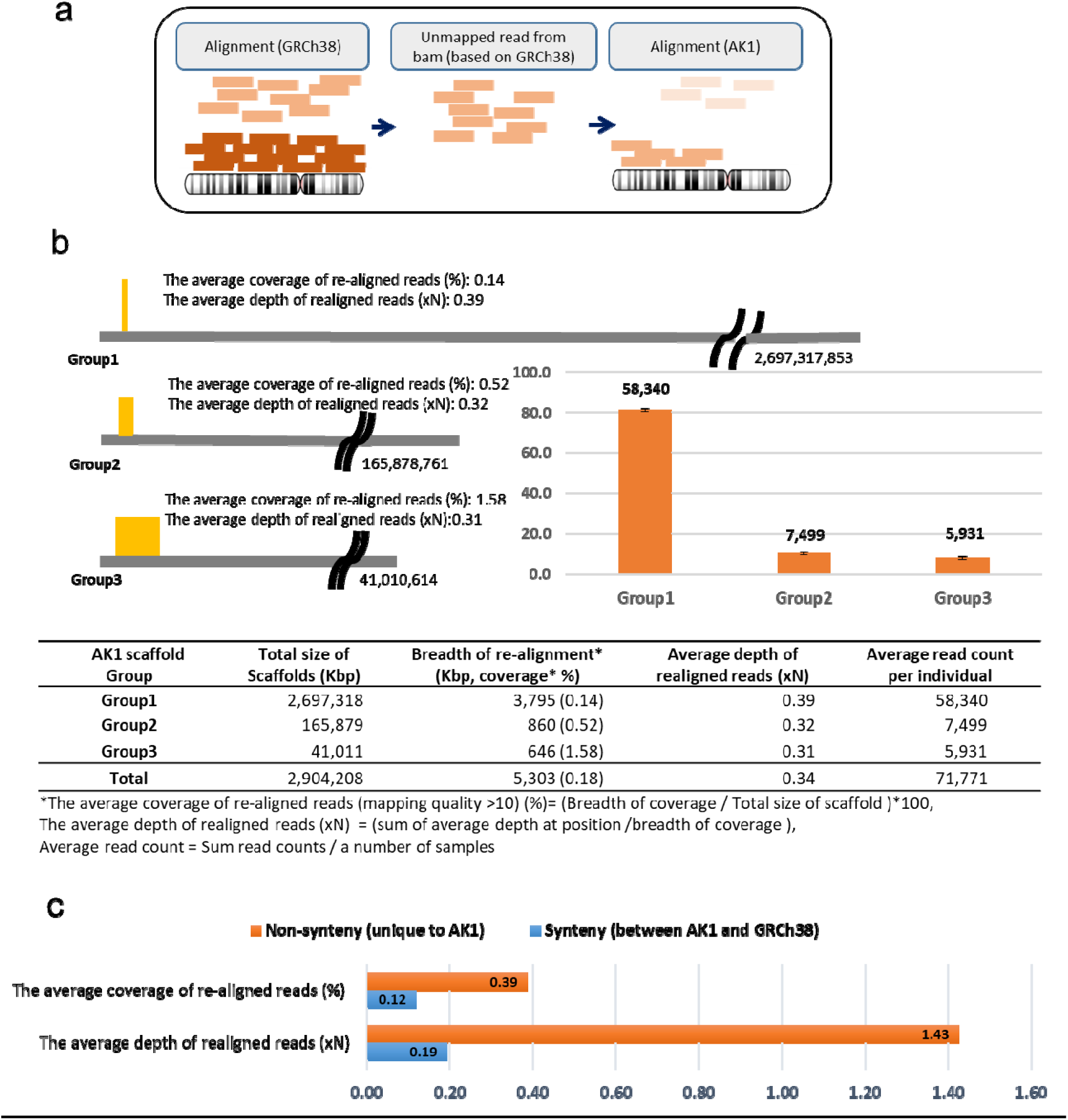
The overall descriptions of “unmapped reads” realigning with AK1. (a) The processes of realigning unmapped reads from GRCh38 reference (b) The breadth of coverage, average depth of coverage by position and read counts by groups. The width of yellow box shows the breadth of coverage and The height of yellow box shows the average depth of coverage by groups. The bar graph(orange) represents a percent of read counts by groups and the numbers above orange bar mean a number of read counts. Error bars represent s.e.m. (c) The average depth and coverage of the recovered sequences of group1 AK1 scaffolds by the synteny status from 14 individuals.

The average coverage of re-aligned reads (%)= (Breadth of coverage / Total size of synteny)*100, The average depth of realigned reads (xN) = (sum of average depth at position /breadth of coverage)

### Characterization of common missing parts

We scrutinized the recovered sequences searching for common missing. We identified 110 regions which were shared by ≥2 individuals with depth >10X for each (Supplemental Table S6). 64 of 110 regions were previously reported or had homology in the BLASTn search of mammalian genome database(Fan et al. 2017; Kehr et al. 2017; Wong et al. 2018)(Supplemental Table S7). Notably, 25 regions showed putative mammalian protein coding functions in the translated BLAST search on NCBI’s nr database (e-value <10^−10^, identity ≥70%, and alignment length ≥50bp). The list of common missing was presented in Supplemental Table S8. We identified that 38 regions were shared globally (≥7 individuals, with depth >10X for each). One of 38 regions was both global missing and suggestive functional, highly homologous with the *zinc finger protein 454 isoform 2* (Supplemental Table S9).

The Group 1 scaffolds of Figure 1 harbored 31 out of 38 globally missing regions; with flanking sequences annotated, the 31 regions were visualized against GRCh38. Typically, those regions were flanked by several repeats elements such as *Alu* or *LINEs* (Supplemental Fig. S3). Further examination of the breakpoints using BioEdit(TA 1999), suggested that non-homologous end-joining with micro-homology (n=26, ∼90%) were the dominant mechanism followed by non-allelic homologous recombination (n=3)(Supplemental Table S10). Interestingly, 26 of the 31 putative insertions had exact matches for chimpanzee, with similar findings for gorilla reference genome (Supplemental Table S10).

We verified the presence of above-mentioned 31 regions which are located within known locations of the reference genome (Group1 of Fig. 1), We conducted PCR amplification using the DNA of AK1, four Europeans and chimpanzee. For AK1, 20 out of 31 putative insertions were verified and 9 regions were also verified for the chimpanzee. For some regions Europeans were either of homozygous or heterozygous for insertion/deletion (Fig. 3).

**Figure 3.**
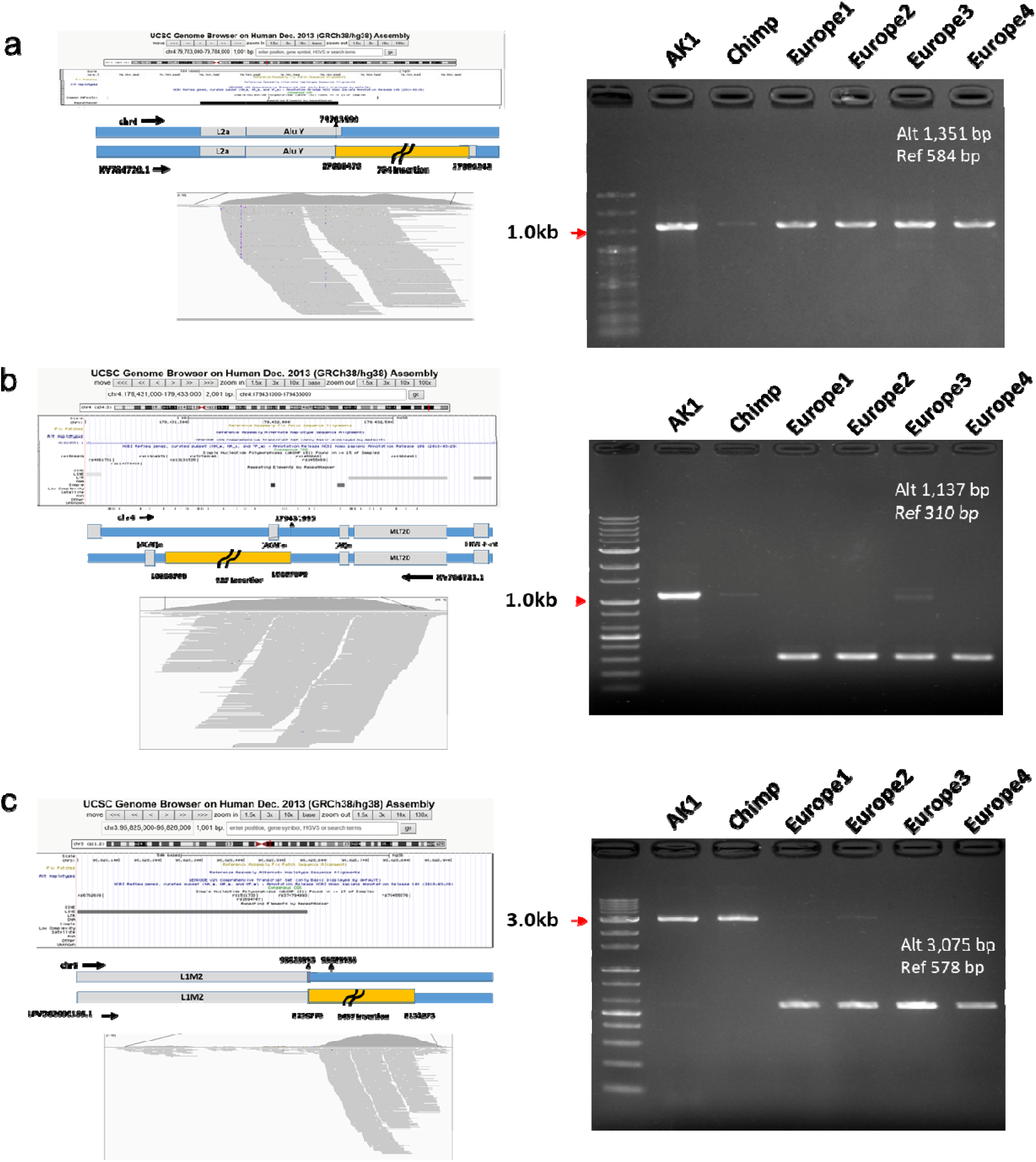
The examples of globally missed regions on GRCh38 investigating with UCSC Genome browser and the experimental verification on the existence of the regions. The regions (Group1) having high depth with 7 or more samples was discovered on the inserted sequences. (a) The region was inserted in AluY (chr4:79,781,761-79,785,451(G1-5)). (b) The region was near the repetitive sequences (chr4:179,430,209-179,433,860(G1-7)) (c) The region was near L1M2(chr3:95,822,539-95,830,080(G1-26)) The yellow block is the inserted sequence against GRCh38, The grey block is repetitive sequences.

## DISCUSSION

By comparison between reference genome and precise ethnic genome, our study suggests that genomic differences between individuals exceed the previous consensus of “99.9% sharing”, which was primarily derived from human genome variation projects; and much less than the 10% of difference derived from the assembly of unmapped reads for African ancestry(Sherman et al. 2018). The magnitude of difference of ∼1.8% might have been either conservative or inflated; it is conservative considering that GRCh38 is a composite genome from the contribution of >50 individuals, and that structural variation was not considered; it might be overestimated in that some of the satellite sequences might not fully identified in both assemblies and those sequences bear unknown functional significances. It is not likely, that both possibilities have substantially affected the estimation of 1.8% difference, considering the level of the accuracy of two genome assemblies. Our findings that ∼0.76% of chromosome is located and arranged differently has less uncertainty, and has significant implications in a range of genome research. In contrast to the estimated difference, only a part of “missing information” was recovered from the unmapped reads (<0.2% of AK1 sequences). It is likely that the differences are attributable to the high proportion of repetitive sequences in unique AK1 regions and to the intrinsic limitations of the analytic platform, e.g., massive short reads of low complexity characterized by the unmapped reads. Our study also discovered that some part of the missing regions are globally common and might harbor functional significances. According to our research about the characterization of common missing parts, the majority of the “globally common missing” regions found from unmapped reads of various populations might be recent deletions in the reference, rather than insertions in other populations. The functional search on globally missing regions of this study was preliminary but limited to coding sequences, so that suggestive functional candidates require further validation.

In conclusion, our study corroborates the usefulness of ethnic genome in acquiring missing genomic information. Highly contiguous ethnic genomes, in particular, will become more attainable, and bear greater importance in understanding complete genome function, as well as more precise human evolutionary history. The precise ethnic genome will also play an important role to redress the large research gap between populations.

## METHODS

### Comparison between the reference genome (GRCh38) and the AK1

We used LASTZ program(Harris 2007) for generating a chain file between the AK1 and GRCh38, with written parameters (-gapped -gap = 600,150,-hspthresh = 4500,-seed = 12of19 –notransition -ydrop= 15000)(Seo et al. 2016). We used UCSC Kent utilities(https://github.com/ENCODEDCC/kentUtils) for the chaining and netting process. Based on synteny and gaps of the chain file, we calculated the alignments between AK1 scaffolds and GRCh38 chromosomes.

### Study samples and materials for profiling mapped/unmapped reads to the reference genome (GRCh38)

The data of all samples were downloaded from the 1000 genome browser and aligned to GRCh38 full analysis set with HLA sequences. All the WGS data were generated by Illumina Hiseq platforms with PCR-free procedures (Genomes Project et al. 2015). We only selected samples of 3 ethnic groups which were deeply sequenced (depth >50X), mapped to GRCh38 with BWA-MEM(version bwakit-0.7.12.)(Li and Durbin 2009), and performed written QC process (ftp://ftp.1000genomes.ebi.ac.uk/vol1/ftp/data_collections/1000_genomes_project) such as sorting, marking duplicates, and Indel realignment by Samtools (version 1.2), BioBamBam (version 0.0.191) (Tischler G 2014;9:13), GATK-3.3-0 (McKenna et al. 2010) and Cramtools.3.0. Resulting 14 individuals ethnicity and race were presented in Supplemental Table S3.

### Investigating features of mapped/unmapped reads from bam files, and re-aligning the extracted “unmapped reads” to the AK1

For qulity check and repetitive annotation of mapped/unmapped reads, we used FastQC(Andrews 2010) and RepeatMasker(Smit 2015). After we extracted unmapped reads from the downloaded bam files of 14 multi-ethnic samples, we realigned unmapped reads to Ak1 genome with using BWA-MEM (Figure 2a). After realigning, the procedures of sorting and removing duplicates were performed by Samtools (version1.3) and Picard-tools(version 2.0.1). We excluded regions with low depth(<3X) for each individual. We handled data and calculated the depth/breadth of re-alignments(Sims et al. 2014) using Bedtools(version 2.25.0)(Quinlan and Hall 2010), Samtools (version1.3) and R(version 3.4.3). We also used GATK-pathSeq (Walker et al. 2018) to identify those reads with the putative microbial sequence.

### Functional search for unique regions of AK1 and the common/globally missing regions

For the sequences (>200bp) unique to the AK1 when we compared between GRCh38 and AK1, we searched for functional clues using a translated BLAST search(BLASTx)(Camacho et al. 2009). Also, to examined whether common missing regions existed across populations and what the functional role of common missing regions which defined as regions mapped from ≥2 individuals with >10 depth, we used both BLASTn and BLASTx search on nr database with default options. The results of BLASTn were filtered with e-value <10^−10^, identity ≥70%, and covering ≥70% and those of BLASTx were filtered with e-value <10^−10^, identity≥ 70%, and alignment length ≥50bp. For further investigation on the globally missing regions, which defined as the regions of common missing regions that shared by seven or more individuals, located in Group1 scaffolds, we searched locations of missing regions on GRCh38 genome using chain file (“lifting” AK1 over GRCh38). To visualize these regions, we merged 14 bamfiles into one and used the UCSC genome browser(Karolchik et al. 2003) and Integrative Genomics Viewer (IGV)(Robinson et al. 2011) on visualization of merged bam file. The suggestive functional role of globally missing regions was also identified with BLASTx search with e-value <10^−10^, identity≥ 70%, and alignment length ≥50bp.

### Verifying the missing regions by PCR

To investigate the characteristics of globally missing regions on Group1 scaffolds, we selected ∼2 kb of flanking sequences on both streams of AK1 and put to BLAST-Like Alignment Tool (BLAT) (http://genome.ucsc.edu/cgi-bin/hgBlat) for human (GRCh38; Dec 2013) and chimpanzee (panTro6; Feb 2018) genome. For experimental confirmation of the non-overlapping AK1 sequences, we performed PCR amplification using four European DNA samples (NA17001, NA17002, NA17003, and NA17004) and a chimpanzee DNA sample which were distributed by the Coriell Institute (Coriell Cell Repository, USA) and provided by Dr. Takenaka (Primate Research Institute, Kyoto University, Japan). Oligonucleotide primers for PCR amplification of each locus were designed by using the software Primer3 (Untergasser et al. 2012). PCR amplification of each locus was conducted with 25 ul reaction using 100 ng DNA, 10μL of 2X Lamp *Pfu* DNA polymerase (BioFact, South Korea), and each oligonucleotide primer (10pmol/μL). The PCR conditions were as follows: 95°C for 5min, followed by 35 cycles of 30 sec of denaturation at 95°C, 40 sec of annealing temperature, and 1 to 7 min of extension at 72°C (depending on the expected size of PCR product), followed by 5 min of a final extension at 72°C.

## DATA ACCESS

When we compared between GRCh 38.p12 and AK1 genome, the chain file was used and supported the findings of this study. The chain file between GRCh 38.p12 and AK1 genome can be found in http://referencegenome.snu.ac.kr/download

The URLs for Ak1 genome and GRCh38 p.12 fasta are each https://www.ncbi.nlm.nih.gov/assembly/GCA_001750385.2 and https://www.ncbi.nlm.nih.gov/assembly/GCA_000001405.27

The downloaded bam files with high coverage are available at ftp://ftp.1000genomes.ebi.ac.uk/vol1/ftp/data_collections/1000_genomes_project

## ACKNOWLEDGMENTS

This research was supported by the Post-Genome Technology Development Program (10050164, Developing Korean Reference Genome) funded by the Ministry of Trade, Industry and Energy (MOTIE, Korea) and the Basic Science Research Program through the National Research Foundation of Korea(NRF) funded by the Ministry of Education(NO. 2019R1A6A3A13093761)

## Author Contributions

J.N.K and J.S. designed the study or supervised analyses. J.N.K, J.S., and K.H. wrote the manuscript. J.N.K, W.S.L., S.Y.M, and J.Y.L. analysed data. All authors reviewed and approved the final manuscript.

## Ethics approval and consent to participate

Not applicable.

## Consent for publication

Not applicable.

